# How small is too small? Genetic signatures from naturally small populations of a large island vertebrate, the Turks and Caicos Rock Iguana

**DOI:** 10.64898/2026.09.13.751268

**Authors:** Zachary Dykema, Glenn P. Gerber, Giuliano Colosimo, Mark E. Welch

**Affiliations:** Department of Biological Sciences, Mississippi State University, Mississippi State, MS, USA; Island Wildlife Alliance, St. Augustine, FL, USA; Department of Biology, University of Rome Tor Vergata, Rome, Italy

**Keywords:** extinction vortex, genetic diversity, population structure, genetic drift, Cyclura, microsatellites

## Abstract

Small populations are increasingly common on the landscape, and many face rising probabilities of extinction. Demographic factors alone can drive a small population into an extinction vortex, but genetic factors such as inbreeding and genetic drift lead to reductions in genetic diversity and expression of deleterious alleles, further impacting fitness. Populations on islands are also impacted by founder effects, minimal gene flow, and habitat constraints, yet naturally small, island populations have persisted through many generations despite all these factors. This suggests that island biota may operate at smaller viable population sizes than their mainland counterparts while avoiding extinction vortices. Using 26 microsatellites, we describe the genetic diversity and population structure of 13 subpopulations of a large island vertebrate, the endangered Turks and Caicos rock iguana (*Cyclura carinata*), found on a range of small cays (0.151–374ha). Most subpopulations had decreased genetic variation but limited evidence of inbreeding despite small island size. Even with water barriers limiting gene flow, some subpopulations in geographical proximity are functioning as a metapopulation while others are not. These subpopulations are not in danger of extirpation, even though census size could be as low as 10 individuals. We believe this study shows that *C. carinata* has a low population size threshold for long-term persistence that can inform the species’ future management, such as with conservation translocations. Furthermore, this project anecdotally adds more evidence to how island populations persist through selection despite genetic drift.

## INTRODUCTION

A human-driven mass extinction event is underway, and habitat loss is a leading factor driving the process (Loarie et al. 2009; Ceballos et al. 2015). Many threatened and endangered species persisting in restricted and fragmented habitats are likely to succumb to extirpation. Demographic and genetic factors, or synergistic effects of the two combined (Gilpin and Soulé 1986), are the anticipated causes leading to further losses, implying that a hefty extinction debt has yet to be realized. Given this likelihood, devising strategies for preserving species with dwindling population sizes has been a high priority for conservation biologists. One of the most pressing questions in this regard is how small is too small?

This simple question has not proven easy to answer, nor has a broad consensus been reached. In the realm of conservation biology, this topic was taken up in earnest in the mid-1970s by wildlife managers concerned with establishing and maintaining “viable populations”, thus preventing extinction (Gilpin and Soulé 1986). Populations face direct, systemic pressures that lead to extirpation (deterministic extinction) as well as stochastic pressures via demography, the environment, genetics, and catastrophes (stochastic extinction) (Shaffer 1981). These drivers are often not mutually exclusive and may ultimately lead to a positive feedback loop, known as an extinction vortex (Gilpin and Soulé 1986). In theory, the pace of extinction vortices should quicken as populations grow smaller because impacts of demographic and environmental stochasticity are exacerbated. Variance in the ability to find mates, reproductive output, and survival rates increase a population’s extinction risk. These compounded effects decrease population fitness and may facilitate population crashes.

Determining a minimum viable population size depends heavily on species, life history, and past population dynamics (Chaudhary and Oli 2020). However, the general 50/500 rule, referring to an effective population size >50 to limit inbreeding depression for at least five generations and an effective population size >500 to limit rates of genetic drift allowing for the retention of evolutionary potential for the long-term, provides guidance for the maintenance of endangered species (Franklin 1980; Soulé 1980). This framework has been further refined and updated (eg. Frankham et al. 2014), and the rule’s usefulness has been both supported and debated (Jamieson and Allendorf 2012; Frankham et al. 2014; Franklin et al. 2014).

Demographics alone in shrinking populations might lead to extinction in the short-term before inbreeding depression has an opportunity to influence the outcome (Melbourne and Hastings 2008). A controversial argument in response to extinction vortices even goes so far as to suggest that extinction almost always occurs too quickly for genetics to play a formative role (Lande 1988). However, the body of scientific literature detailing inbreeding depression’s impact on laboratory and natural populations has confirmed the importance of genetic factors in the calculus of extinction (Frankham 2005; Hasselgren and Norén 2019). Inbreeding and loss of genetic diversity affect extinction risk by reducing reproduction and survival. Though harder to parse out in wild populations due to the difficulty of detecting inbreeding depression without pedigrees (Keller and Waller 2002a), recent studies using genomics continue to provide evidence for decreased fitness as consanguineous matings increase (Kardos et al. 2016). Adaptative potential in the face of environmental changes (e.g. new diseases, predators, or climate change) is impacted by decreased genetic diversity in small populations (Frankham and Kingsolver 2004). In fact, increasing genetic diversity via translocations (i.e. genetic rescue) in inbred, natural populations has been inferred to have successfully reverted extinction vortices in Florida panthers (Pimm et al. 2006), greater prairie chickens (Westemeier et al. 1998), and European adders (Madsen et al. 2004), demonstrating that population genetics can be used to manage extinction risk (but see Hedrick et al. 2014).

A key concern from a population genetics perspective when considering population viability is the enhanced role that genetic drift should play in small populations. Large populations are less impacted by genetic drift because there are more mating opportunities (Rich et al. 1979; Lohr et al. 2014). This implies that rare beneficial alleles are more likely to be retained since there is effectively a large sample size of alleles being passed from one generation to the next. Small populations, on the other hand, provide fewer mating opportunities, causing allele frequencies between parent and offspring generations to be less similar due to chance. Rare alleles are more likely to be lost, and common alleles are more likely to become fixed. Hence, small populations impacted by genetic drift should be more homozygous and maintain fewer rare alleles than their larger counterparts. In theory, the frequency of some deleterious alleles should also increase due to genetic drift. While natural selection should filter out less fit individuals and purge some portion of the genetic load, purging may be insufficient to maintain population fitness (Reed and Bryant 2001; Reed et al. 2003). Selection acting against one component of genetic load could facilitate the fixation of others, furthering the effects of inbreeding depression (Whitlock 2000).

Island biotas have been informative when evaluating the limits of population size (MacArthur and Wilson 1967; Hanski and Kuitunen 1986; Gao and Wang 2022). Even with restricted habitat and isolation that heighten extinction risk compared to mainland populations, island species manage to persist with low census population sizes and often undergo remarkable radiations over time (Frankham 1998; Losos and Ricklefs 2009). Metapopulation processes such as migration, extirpation, colonization, and local selection all shape island populations (Hanski 1998). Many island species are believed to have been established with few founders, implying they passed through genetic bottlenecks, yet they clearly overcame limited population size and genetic diversity (Frankham 1997, 1998). Island carrying capacities are constrained by their size and limited gene flow due to water barriers (Richman et al. 1988). These further limit population growth and genetic variability over many generations, thus reducing effective population size, hindering adaptability, and increasing extinction risk (but see James et al. 2016).

Islands formed by rising sea level following the last glacial maximum should be of special interest. Extant populations residing on small islands formed by rising sea levels 20,000–3,000 years ago have endured habitat loss, reduced connectivity, and elevated levels of inbreeding (Frankham 1997). Furthermore, these island populations have also clearly maintained sufficient genetic variation to adapt to the changes in climate accompanying the release of the glacial cycle (e.g. Cunha et al. 2011). Hence, these archipelagic species can be viewed as survivors of a process currently being endured by mainland taxa facing habitat loss, fragmentation, and climate change.

The Turks and Caicos Islands (TCI) and the Turks and Caicos rock iguana (*Cyclura carinata*) provide an excellent opportunity to explore genetic conditions under which a terrestrial vertebrate species might survive in the long-term following massive habitat loss. This archipelagic region is made up of over 200 islands ranging from 0.10 to 13,500 hectares (Fig. 1). It should thus be feasible to conduct comparative analyses of genetic diversity among island populations of varying sizes. However, many of the islands on the Caicos Bank are only isolated by shallow water channels and undergo periods of connectivity via sand banks that form due to tides, currents, and storms (Welch et al. 2004). It seems likely that at least some of these clusters of islands function as metapopulations for local flora and fauna. Bathymetry implies that other TCI cays have been completely isolated for thousands of years (Lighty et al. 1982; Sealey and Logan 2013).

**Figure 1:**
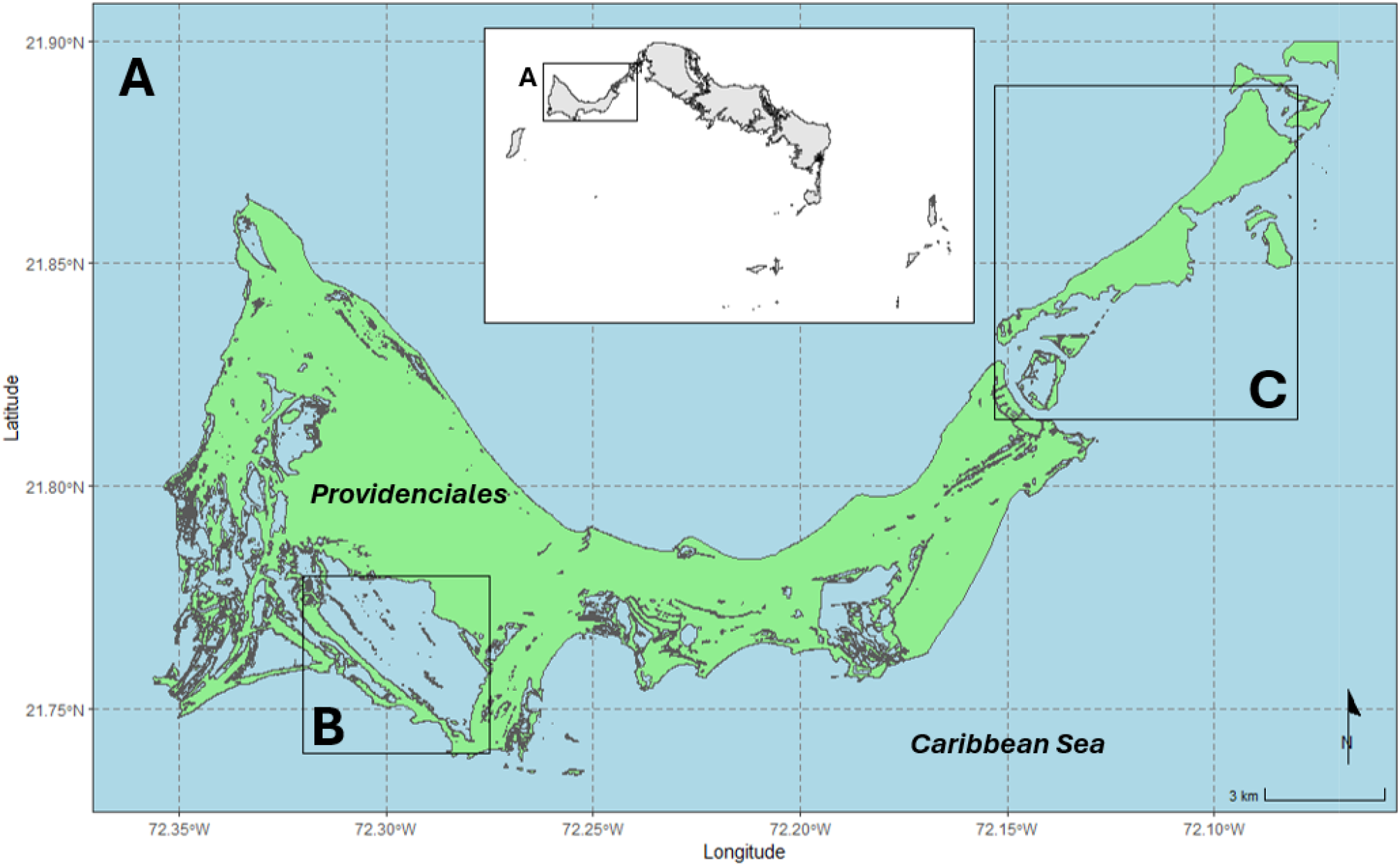
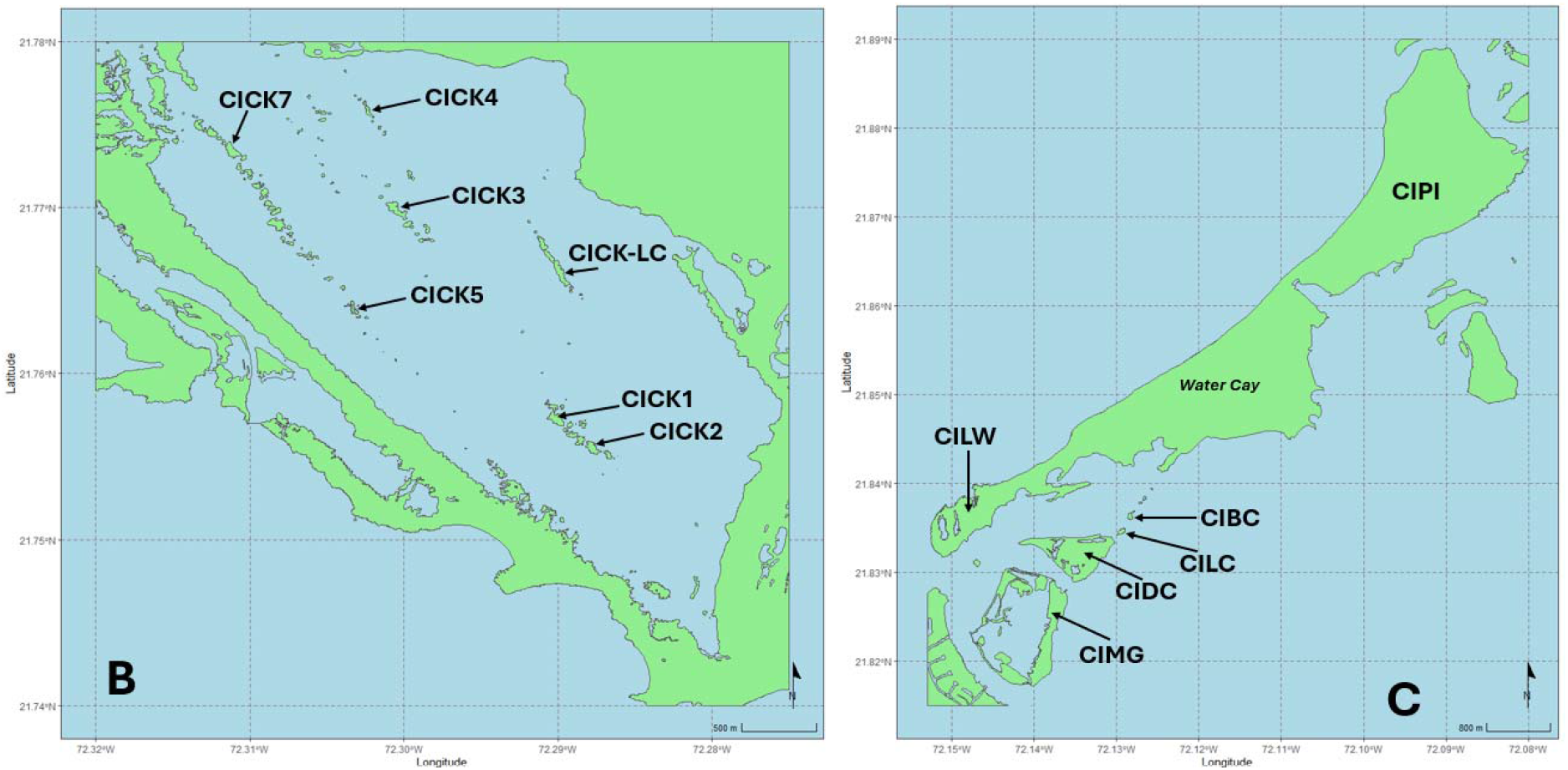
A) Map of sampling locations surrounding Providenciales in the Turks and Caicos Islands. Inset map is the entire Turks and Caicos island chain. B) Sampling locations in Chalk Sound National Park. C) Sampling locations of outlying cays including Little Water Cay, an iguana sanctuary. Water Cay was not sampled for this project. [**Alt text:** Map of the Turks and Caicos Islands with two panels zoomed in on the two study regions: Chalk Sound National Park and the northeast outlying cays.]

**Figure 2:**
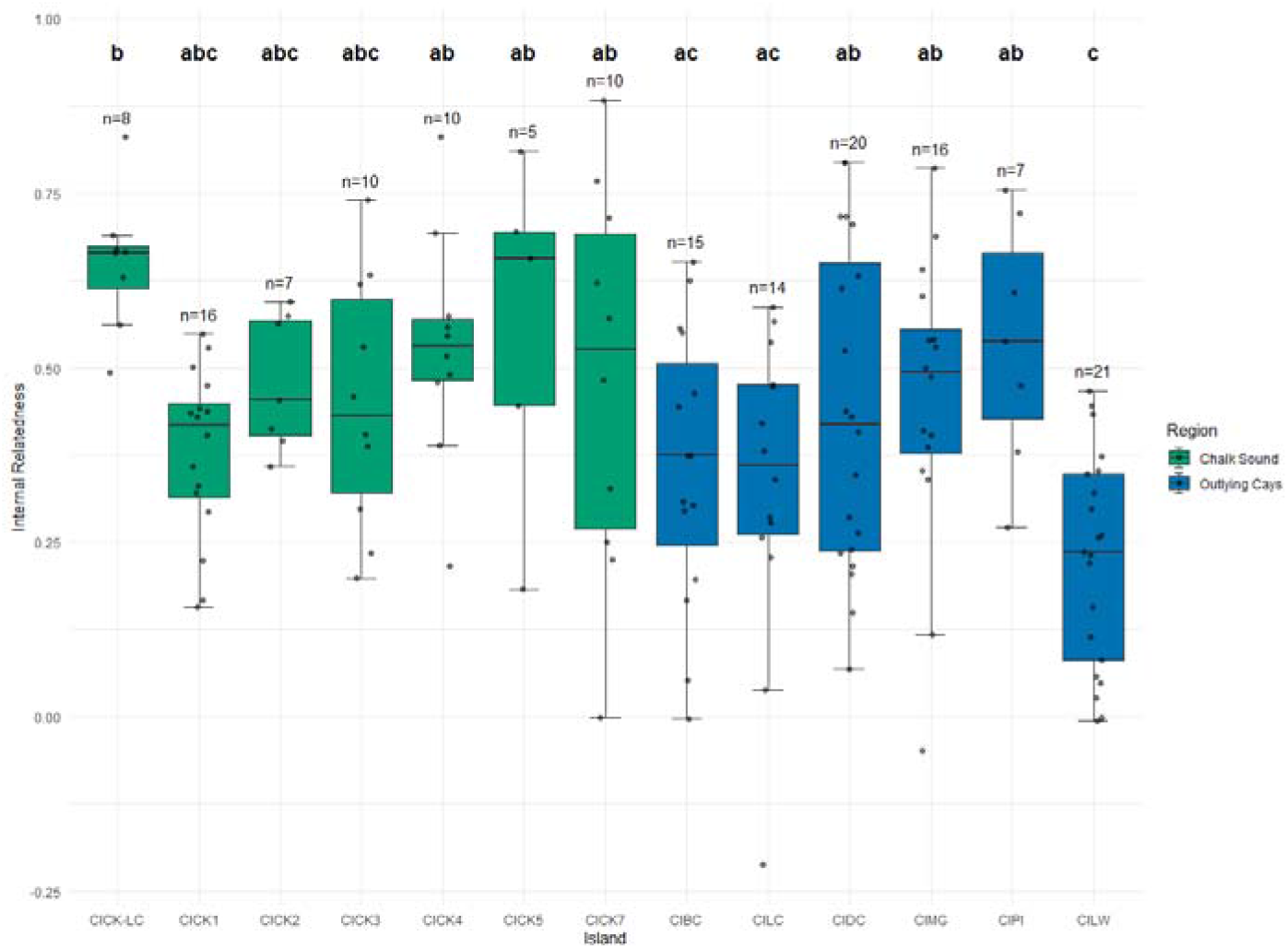
Boxplots of internal relatedness values for each iguana on all 13 sampled cays, color-coded by region. Sample size (n) is above each boxplot. Plots that do not share a letter are significantly different from each other based on an ANOVA and Tukey HSD test. [**Alt text:** Boxplots comparing internal relatedness across 13 subpopulations with statistical differences indicated over each plot.]

*C. carinata* is the largest, extant, native terrestrial vertebrate endemic to the TCI, and it is classified as endangered by the IUCN Red List (Gerber et al. 2020). It is estimated that the historic range of *C. carinata* has decreased by more than 90% due to anthropogenic impacts (Gerber et al. 2020). Rock iguanas are important seed dispersers, browsers, and soil cyclers (Iverson 1985). The total census population size of *C. carinata* is currently estimated to be around 30,000 individuals. However, the population is fragmented across dozens of islands and operates as multiple, naturally small subpopulations limited by island carrying capacity and the presence of invasive alien species limiting the potential for expansion and colonization (Welch et al. 2017; Colosimo et al. in review).

In this study, we used microsatellites to compare genetic variability among multiple small subpopulations of *Cyclura carinata* that vary in population size. Assuming these subpopulations have remained mostly isolated since the Last Glacial Maximum, their persistence suggests that they are sufficiently large to maintain long-term viability. This system therefore provides an opportunity to examine how population size influences genetic diversity and to evaluate alternative mechanisms that may mitigate the effects of genetic drift. We explored three different explanations of heterozygosity in this system. First, if genetic drift is the primary force shaping genetic diversity, we predicted that heterozygosity would decrease with decreasing population size. Second, if processes other than genetic drift contribute to the maintenance of genetic diversity, such as selection, we predicted that heterozygosity would be greater than expected from genetic drift alone. Third, if some subpopulations are not genetically independent but instead function as components of larger metapopulations, we predicted that gene flow among subpopulations would mitigate the loss of heterozygosity expected from drift. Together, these analyses provide empirical insight into the population size required to maintain genetic diversity in *C. carinata* and may inform the management of small populations, including those established through conservation translocations.

## METHODS

### Study system

*Cyclura carinata* has a polygynous mating system, and both sexes compete for territories containing nesting habitat, resources, and mates (Iverson 1979, Gerber et al. 2020). Hence, island size, and by proxy amount of suitable habitat, is expected to significantly limit iguana population size. Iguana-inhabited islands in the TCI range widely in size, from 0.151–440ha. Across the species range, maximum population density is estimated to be ∼30 adults per hectare or ∼60 total individuals across age classes per hectare (Gerber 2004). Nevertheless, multiple cays in the western portion of the Caicos Bank, though <1.0ha, have sustained iguana populations. This study focuses on the western Evolutionary Significant Unit (ESU) described in previous studies, which encapsulates iguanas found in Chalk Sound National Park, Providenciales, and six cays proximal to Providenciales (Fig. 1A, Welch et al. 2017). Researchers believe these naturally small subpopulations result from isolation associated with the rising sea levels of the current interglacial (Welch 1997, Welch et al. 2004, Welch et al. 2017). Increased sample size and number of cays sampled allow for the exploration of the fine-scale population structure of this ESU.

Chalk Sound National Park is located on the southwestern coast of Providenciales (Fig. 1B), one of the larger islands of the Caicos Bank (∼10,500ha). Established in the late 1980s under the jurisdiction of the Turks and Caicos Department of Environment and Coastal Resources, Chalk Sound National Park consists of dozens of small, vegetated, limestone cays surrounded by a shallow (∼1.5m) lagoon (Mitchell and Barborak 1991). Some of these cays have subpopulations of *C. carinata* isolated from invasive mammals due to water barriers. Iguanas in Chalk Sound National Park are presumed to be remnants of the extirpated subpopulation once found on the rest of Providenciales.

Similarly sized cays with iguana populations are also found on the northeast side of Providenciales in the Princess Alexandra Nature Reserve (Fig. 1C). Deep water channels and powerful tide dynamics isolate Bird Cay, Lizard Cay, Donna Cay, and Mangrove Cay from neighboring islands, including Little Water Cay, Water Cay, and Pine Cay, which are currently connected by sand bars forming a single landmass. Feral cats were present on this interconnected landmass, mostly on Water Cay and Pine Cay, until an eradication program took place in 2017 and 2019. In 2020, when Pine Cay was sampled for this study, the iguana populations on Water Cay and Pine Cay were in the early stages of population rebound. Except for Water Cay and Pine Cay, these outlying islands have been largely buffered from human impacts. Thus, their respective iguana populations have persisted, at or close to carrying capacity, while iguana populations on larger, developed islands declined or were extirpated.

Little Water Cay (CILW) is a government-protected, ∼40-hectare National Park with one of the densest iguana populations in *C. carinata*’s native range, and census population estimates are ∼2000 individuals (Gerber 2007). Geological and genetic studies indicate that the CILW subpopulation has largely been isolated from other cays, though intermittent connections to other islands, such as the sandbars now connecting CILW to Water Cay and Water Cay to Pine Cay, have likely provided periodic opportunities for gene flow (Fig. 1A, Gerber 2007). All sampled subpopulations are listed in Table 1 and mapped in Figure 1.

**Table 1:** List of 13 *C. carinata* subpopulations sampled in this study with island region, island name, island name code, island area in hectares, approximate census size, and number of blood samples genotyped. Asterisks indicate cays assumed to be at carrying capacity based on previous studies.

| Region | Island | Code | Area (ha) | Sample size |
| --- | --- | --- | --- | --- |
| Chalk Sound National Park | Chalk Sound Long Cay | CICK-LC | 0.86 | 8 |
|  | Chalk Sound 1 | CICK1 | 0.71 | 16 |
|  | Chalk Sound 2 | CICK2 | 0.16 | 7 |
|  | Chalk Sound 3 | CICK3 | 0.65 | 10 |
|  | Chalk Sound 4 | CICK4 | 0.23 | 10 |
|  | Chalk Sound 5 | CICK5 | 0.15 | 5 |
|  | Chalk Sound 7 | CICK7 | 0.53 | 10 |
| Northeast outlying cays | Bird Cay | CIBC | 0.42 | 15 |
|  | Lizard Cay | CILC | 0.44 | 14 |
|  | Donna Cay | CIDC | 35.2 | 20 |
|  | Mangrove Cay | CIMG | 116.0 | 16 |
|  | Pine Cay | CIPI | 374.0 | 7 |
|  | Little Water Cay* | CILW | 34.9 | 21 |

### Data Collection and Genotyping

Sampling took place across 12 cays (see Table 1) during the summers of 1995 and 1997, and Pine Cay during the spring of 2020. Adult iguanas were captured by hand or noose pole and individually marked with colored beads to avoid recapture. Blood was collected (0.2-1.0 mL) via the ventral coccygeal vein using heparinized syringes and stored in SDS lysis blood buffer (0.1M Tris-HCl pH 8.0, 0.1M EDTA, 0.01M NaCl, 0.5% SDS) at ambient temperature in the field, and later at -80°C for long-term storage.

DNA was extracted from blood using a Promega Maxwell® 16 and proprietary chemistry (Promega, Madison, WI, USA). Microsatellites known to be polymorphic in *C. carinata* (Welch et al. 2017, Table S1) were PCR amplified and screened for variability in the iguanas sampled for this study using previously described methods (Welch et al. 2011, 2017). Fragment analysis was performed on 3730xl DNA Analyzers (Applied BioSystems, Foster City, CA, USA) at the Cornell Institute of Biotechnology Core Facility using LIZ-500 as size standard (GeneScan 500 LIZ Size Standard, Applied BioSystems). Genotypes were scored using PEAK SCANNER version 1.0 (Applied BioSystems).

### Estimation of Summary Statistics

Loci found to be invariant were omitted from further analysis. To ensure measures of genetic diversity were robust, genotypes generated for each locus in each subpopulation were tested for the presence of null alleles and large allele dropout using FreeNA (Chapuis and Estoup 2007; Chapuis et al. 2008). This software estimates null allele frequencies using the Expectation Maximization (EM) algorithm (Dempster et al. 1977).

Unless otherwise specified, R version 4.3.2 (R Core Team 2019) was used for the remaining analyses. Departures from Hardy-Weinberg equilibrium (HWE) were assessed for statistical significance using *hw.test()* in the R package pegas (Paradis 2010). Specifically, we conducted 1000 Monte Carlo permutations of the exact test (Guo and Thompson 1992) while controlling the false discovery rate of our p-values using the base stats *p.adjust()* (Benjamini and Hochberg 1995). Besides total count of alleles per subpopulation sampled, we also calculated rarefied allelic richness (Ar) using *allel.rich()* in package PopGenReport (Adamack and Gruber 2014). This rarefaction method uses the smallest number of alleles in a sample (in our case, 2) as the sample size (El Mousadik and Petit 1996). Count of private alleles unbiased for sample size was calculated for each subpopulation using the formula *A*′ = *A* × (*n’* / *n*), where *A* is the number of private alleles in a subpopulation, *n’* is the minimum sample size out of all subpopulations, and *n* is the actual sample size of the subpopulation (Kalinowski 2004). List and number of private alleles were generated using *private_alleles()* in package poppr (Kamvar et al. 2014). Observed heterozygosity (*H_O_*), expected heterozygosity (*H_E_*), and *F_IS_* were calculated using formulas in Weir and Cockerham 1984 through the *summary()* command in package adegenet (Jombart 2008).

Multi-locus heterozygosity (MLH) within individuals across sampled cays was estimated and differences were tested for statistical significance. Internal relatedness (IR) values range from -1 for individuals that are maximally outbred to 1 for individuals that are maximally inbred, based on relative levels of multi-locus heterozygosity given allele frequencies within populations (Amos et al. 2001). IR was deemed the most appropriate measure of MLH because it operates well when average heterozygosity is low (Aparicio et al. 2006). IR was calculated for all sampled individuals using the *mlh()* command in the package Rhh v. 1.0.2 (Alho et al. 2010). We conducted an ANOVA and Tukey HSD test to detect significant differences in IR between subpopulations.

Iguana population size is in direct proportion to island size. As a proxy for census size, we conducted simple linear regression models between log_10_ transformed island surface area and three genetic metrics: *H_E_*, *F_IS_*, and IR. Island surface area was estimated for each cay using the measuring tool in Google Earth. Base R was used to create the models. Effective population size (*N_E_*) was estimated with NeEstimator V2.1 (Do et al. 2014). The software uses different algorithms to infer contemporary *N_E_*: one based on linkage disequilibrium (Waples and Do 2008), one based on excess heterozygosity (Zhdanova and Pudovkin 2008), and one based on molecular coancestry (Nomura 2008). As a comparison, *N_E_* was also estimated using the drift-mutation equilibrium equation 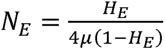 where *H_E_* is Weir and Cockerham’s estimate of expected heterozygosity and is the mutation rate (Crow and Kimura 1970). We used a median human microsatellite mutation rate, 1.8 × 10^−3^, for (Sun et al. 2012).

### Population Structure

Many cays, though separated by water barriers, may be operating as metapopulations and not as independent subpopulations. *Cyclura carinata* avoid water but can swim and will do so when close to land (Iverson 1979, Gerber pers. comm.). Tropical storms and subsequent vegetation rafts formed may play an important role in iguana dispersal (Censky et al. 1998). A Mantel test was conducted to determine whether the correlation among genetic and geographic distances was significant (Mantel 1967), using *mantel.rtest()* in the package ade4 (Bougeard and Dray 2018). Pairwise *F_ST_*values (Weir and Cockerham 1984) were compared using *pairwise.WCfst()* in the package hierfstat (Goudet 2005). To test allele frequency differences for statistical significance, 1000 bootstraps over all loci were run to test if values were significantly different from 0 using *boot.ppfst()*. An AMOVA was also conducted using package pegas with 999 permutations (Paradis 2010).

To further test for population structure among the sampled cays, we conducted a discriminant analysis of principal components (DAPC, Jombart et al. 2010). Retention of principal components was decided based on first running *dapc()* from the package adegenet using n.pca=50, then calculating a-scores for optimization (Jombart et al. 2010). Cross validation was done using *xvalDapc()* in the same package. The command *dapc()* was run again with the optimal number of principal components. The Bayesian individual assignment test implemented in STRUCTURE v 2.3.4 was used as a parallel analysis (Pritchard et al. 2000). We ran the program with a flat prior for the admixed parameter and tested values of K (number of genetic clusters) from 1 to 13 with 20 replicates of each (Gilbert et al. 2012). STRUCTURE ran 10^5^ Markov Chain Monte Carlo (MCMC) as burn-in for each run and another 10^5^ MCMC as replicates. Output from STRUCTURE was uploaded to the web app POPHELPER (Francis 2017) to calculate the second order of differences in the likelihood function of K[ΔK] and create plots (Evanno et al. 2005).

## RESULTS

### Genetic Diversity

In total, 159 iguanas were sampled with the number of samples from each cay ranging from 5 to 21 (median = 10, see Table 1). Two of the 28 microsatellite loci were monomorphic (CIDK109, CIDK135) and were omitted from further analyses (Table S1). We found few instances where estimated null allele frequency was greater than 0.20 across the 26 loci and 13 cays sampled (19/338=0.0562, Fig. S1). After adjusting for the false discovery rate associated with multiple testing, only one locus (D140) deviated from HWE on one cay (CILC, Fig. S2). We retained all remaining 26 loci because significant null allele frequencies and deviations from HWE were negligible.

CILW had the higher number of alleles (Ar = 1.380) compared to Chalk Sound National Park (Mean Ar = 1.229) and the outlying cays (Mean Ar = 1.315), though CIPI had the highest (Ar = 1.404, Table 2). CILW had the highest heterozygosity (*H_O_* = 0.378), followed by the outlying cays (Mean *H_O_* = 0.276), then Chalk Sound National Park (Mean *H_O_* = 0.231). However, both CIPI and CILC had more private alleles when compared to CILW. Most Chalk Sound National Park cays had an inbreeding coefficient indicating moderate heterozygosity excess (Mean *F_IS_* = -0.043) while the outlying cays showed moderate homozygosity excess (Mean *F_IS_* = 0.099). CILW’s inbreeding coefficient was close to 0 (*F_IS_* = -0.007). CIMG and CIPI had an inbreeding coefficient >0.10, indicating some level of inbreeding. Surprisingly, the lowest *F_IS_*was -0.220 in CICK2, showing substantial heterozygote excess.

**Table 2:** Population genetics summary statistics for 13 sampled subpopulations including sample size (n), total number of alleles (A), rarefied allele count adjusted for sample size (Ar), number of private alleles adjusted for sample size (Private), observed heterozygosity (*H_O_*), expected heterozygosity (*H_E_*), and inbreeding coefficient (*F_IS_*). Asterisks indicate cays assumed to be at carrying capacity based on previous studies.

| Region | Island | $n$ | $A$ | $Ar$ | $Private$ | $H_O$ | $H_E$ | $F_{IS}$ |
| --- | --- | --- | --- | --- | --- | --- | --- | --- |
| Chalk Sound National Park | CICK-LC | 8 | 40 | 1.168 | 0.000 | 0.163 | 0.163 | 0.071 |
|  | CICK1 | 16 | 50 | 1.276 | 0.313 | 0.282 | 0.271 | -0.052 |
|  | CICK2 | 7 | 39 | 1.184 | 0.000 | 0.225 | 0.177 | -0.220 |
|  | CICK3 | 10 | 47 | 1.255 | 0.000 | 0.268 | 0.247 | -0.076 |
|  | CICK4 | 10 | 48 | 1.212 | 1.000 | 0.215 | 0.206 | -0.032 |
|  | CICK5 | 5 | 46 | 1.277 | 1.000 | 0.229 | 0.253 | 0.039 |
|  | CICK7 | 10 | 48 | 1.232 | 3.500 | 0.233 | 0.224 | -0.034 |
| Northeast outlying cays | CIBC | 15 | 51 | 1.249 | 0.000 | 0.267 | 0.244 | -0.046 |
|  | CILC | 14 | 55 | 1.283 | 7.143 | 0.297 | 0.278 | 0.012 |
|  | CIDC | 20 | 65 | 1.292 | 1.750 | 0.246 | 0.287 | 0.097 |
|  | CIMG | 16 | 69 | 1.347 | 0.000 | 0.242 | 0.340 | 0.291 |
|  | CIPI | 7 | 59 | 1.404 | 19.286 | 0.327 | 0.383 | 0.142 |
|  | CILW* | 21 | 74 | 1.380 | 4.762 | 0.378 | 0.376 | -0.007 |

We found a significant, positive relationship between log island surface area and *H_E_* as well as log island surface area and *F_IS_*(Fig. 3A, p = 0.0048 and Fig. 3B, p = 0.0004 respectively). No significant relationship was detected between log island surface area and IR, though IR slightly decreased with island size (Fig. 3C, p = 0.6138). Most estimates of effective population size *N_E_* were low (>10), but differences in NeEstimator methodologies and sample sizes led to large confidence intervals (Table 3). *N_E_*estimates using mutation-drift equilibrium were higher than the other estimates, indicating heterozygosity is higher than expected from drift alone (Table 3).

**Figure 3:**
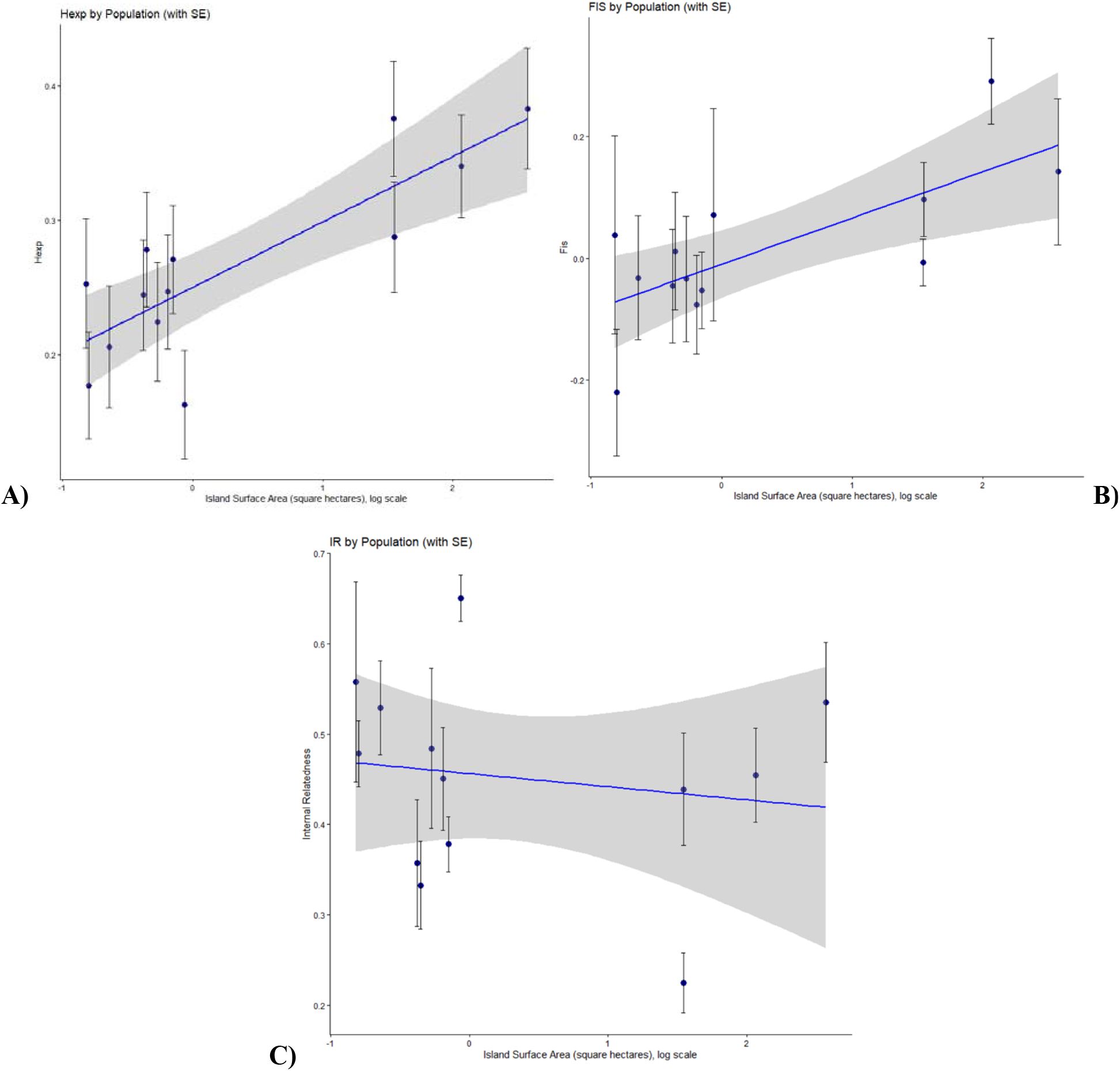
Simple linear regression models for cay surface area on a log scale versus A) *H_E_* (standard error based on number of loci), B) *F_IS_*(standard error based on number of loci), and C) internal relatedness (standard error based on number of samples). Shaded area is the standard error of the model. A) and B) have simple linear models that are significant. [**Alt text:** Graphs comparing log island surface area with changes in expected heterozygosity, inbreeding coefficients, and internal relatedness.]

**Table 3:** One-sample estimates of effective population size (*N_E_*) from the program NeEstimator V2.1. *N_E_* was calculated using the linkage disequilibrium method, heterozygosity excess method, and the molecular coancestry method using number of samples *n*. Least allele frequency (LAF) cutoff was 0.02. Values in parentheses values are 95% confidence intervals of each estimate. *N_E_* was also estimated using Crow and Kimura 1970 mutation-drift equilibrium using island estimates of *H_E_*. Asterisks indicate cays presumed to be at carrying capacity based on previous studies.

| Region | Island | $n$ | Linkage Diseq. | Het. Exc. | Mol. Coanc. | Crow & Kimura |
| --- | --- | --- | --- | --- | --- | --- |
| Chalk Sound National Park | CICK-LC | 8 | 2.9 (1.0-Inf.) | Inf. (5.7-Inf.) | Inf. (Inf.-Inf.) | 27.0 |
|  | CICK1 | 16 | Inf. (20.6-Inf.) | 70.7 (4.9-Inf.) | Inf. (Inf.-Inf.) | 51.6 |
|  | CICK2 | 7 | Inf. (2.0-Inf.) | 5.0 (2.1-Inf.) | 17.9 (0.0-89.9) | 29.9 |
|  | CICK3 | 10 | 5.4 (1.8-68.4) | Inf. (4.0-Inf.) | 217.5 (0.2-1092.1) | 45.5 |
|  | CICK4 | 10 | Inf. (29.7-Inf.) | Inf. (5.4-Inf.) | 4.5 (1.7-8.6) | 36.0 |
|  | CICK5 | 5 | Inf. (1.8-Inf.) | Inf. (Inf.-Inf.) | 6.2 (2.3-12.1) | 47.0 |
|  | CICK7 | 10 | 2.5 (0.8-Inf.) | Inf. (5.3-Inf.) | 1.8 (1.0-3.0) | 40.2 |
| Northeast outlying cays | CIBC | 15 | 3.1 (2.0-8.6) | Inf. (3.8-Inf.) | 3.2 (1.7-5.2) | 44.9 |
|  | CILC | 14 | 1.4 (1.1-1.9) | Inf. (5.5-Inf.) | 1.0 (0.7-1.3) | 53.5 |
|  | CIDC | 20 | 35.4 (14.8-Inf.) | Inf. (Inf.-Inf.) | 2.2 (0.9-4.1) | 56.0 |
|  | CIMG | 16 | Inf. (32.6-Inf.) | Inf. (Inf.-Inf.) | 5.9 (1.4-13.5) | 71.6 |
|  | CIPI | 7 | Inf. (10.7-Inf.) | Inf. (Inf.-Inf.) | 14.0 (1.0-43.8) | 86.3 |
|  | CILW* | 21 | 67.1 (28.6-Inf.) | Inf. (10.7-Inf.) | 7.0 (3.0-12.7) | 83.5 |

### Population Structure

The Mantel test indicates a significant association between geographic and genetic distances among cays, suggesting isolation by distance (Fig. S3A, r = 0.715, p = 0.001). This pattern is not significant among cays in Chalk Sound National Park (Fig. S3B, r = 0.203, p = 0.196) or among the outlying cays (Fig. S3C, r = 0.575, p = 0.100). Most pairwise *F_ST_* values were significant and ranged from 0.111-0.584 (Fig. S4). The only exception was between CIMG and CILW (*F_ST_* = 0.0076). Pairwise *F_ST_* values among cays in Chalk Sound National Park (0.111-0.388) are similar to those among the outlying cays (0.112-0.479). CICK-LC was an outlier amongst the Chalk Sound National Park with pairwise *F_ST_* = 0.331-0.512. The AMOVA indicates genetic structure is significant (F_ST_ = 0.572, p < 0.001) and that allele frequency differences among populations within regions (Chalk Sound and outlying cays) explain a greater proportion of the total genetic variance (F_SC_ = 0.437) than do differences between regions (F_CT_ = 0.239).

DAPC shows groups by geographic region with substantial overlapping in Chalk Sound National Park subpopulation ellipses and separation in the outlying cays (Fig. 4). CIPI separates out as an independent cluster adjacent to the other outlying cays (Fig. 4). The second order of differences in the likelihood function of K[ΔK] indicated K = 2 as the most likely number of genetic clusters represented in the sampled iguana subpopulations, separated by cays in Chalk Sound National Park and all other outlying cays (Table S2, Figure S5). These results, paired with significant IBD between cay regions, led us to repeat DAPC and STRUCTURE separately by region without CIPI.

**Figure 4:**
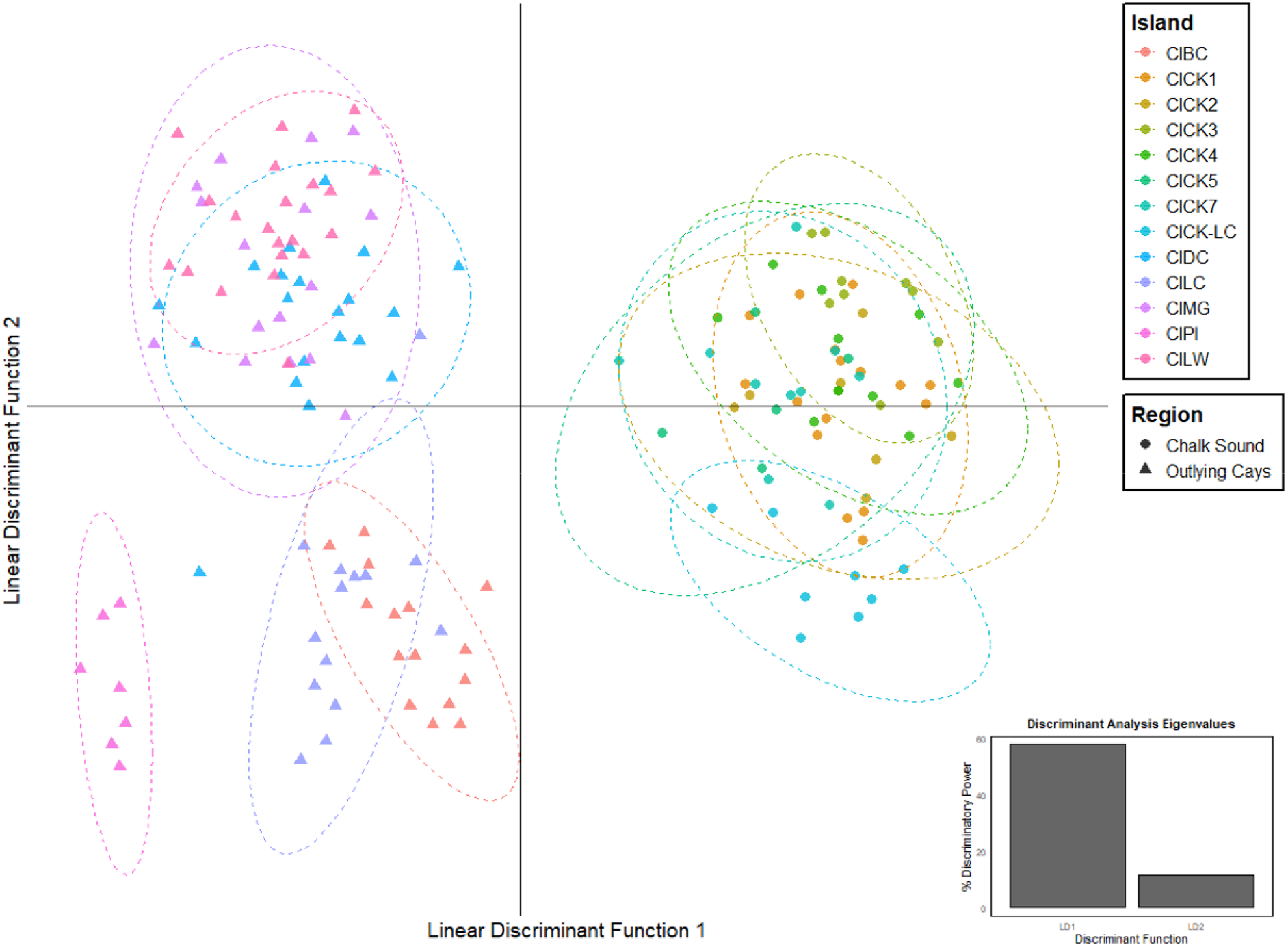
Discriminant analysis of principal components (DAPC) with all sampled cays, coded by sampling location (color) and geographic region (shape). Inset plot is the discriminant analysis eigenvalues of the two principal axes, LD1 and LD2. [**Alt text:** Chart depicting genetic differences in 13 subpopulations, both Chalk Sound National Park and the northeast outlying cays, in discriminant space.]

Chalk Island National Park samples show overlap between ellipses with no island appearing as an isolated cluster (Fig. 5A). Chalk Sound National Park second order of differences in the likelihood function of K[ΔK] indicated K = 3, grouping by geographic distance (Table S3, Fig. S6). However, the change in K[ΔK] going from K = 1-7 changed very little, indicating little genetic structure (Table S3). CIBC and CILC have very little overlap with each other or the rest of the outlying cays (Fig. 5B). Outlying cays second order of differences in the likelihood function of K[ΔK] best support K = 2 with clustering separated by CIBC+CILC and CIDC+CIMG+CILW (Table S4, Fig. S7). There were two individuals of note that were assigned to a different cluster than their sampling location. One CILW iguana had a majority assignment to the CIDC cluster, while another iguana from CICK5 matched very closely with the CICK7 cluster (Fig. 5, Fig. S6E, Fig. S7E).

**Figure 5:**
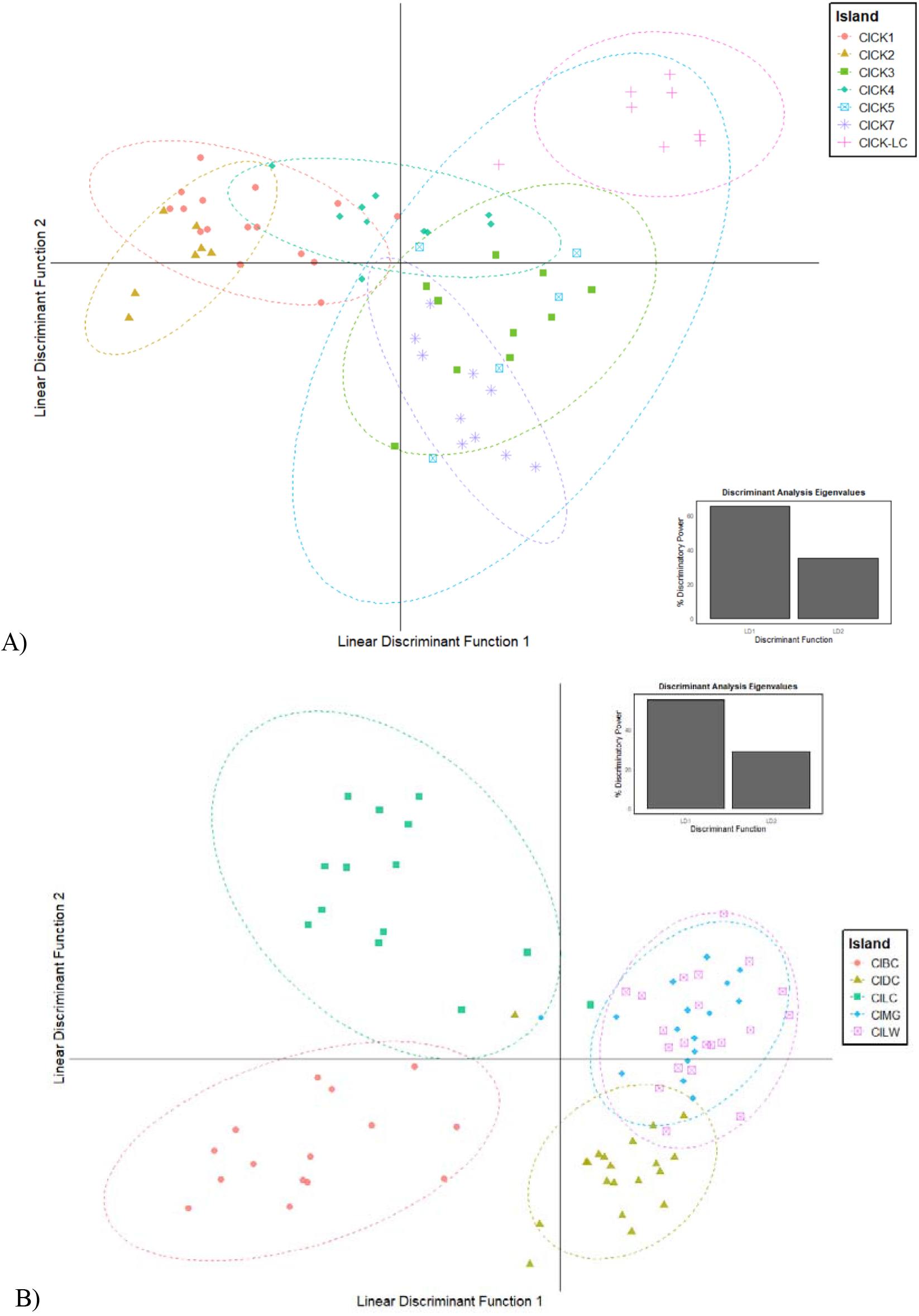
Discriminant analysis of principal components (DAPC) with all sampled cays, coded by sampling location (color) and geographic region (shape). Inset plot is the discriminant analysis eigenvalues of the two principal axes, LD1 and LD2. [**Alt text:** Chart depicting genetic differences in 13 subpopulations, both Chalk Sound National Park and the northeast outlying cays, in discriminant space.]

## DISCUSSION

Small populations of plants (Matthies et al. 2004) and animals (Pimm et al. 1993) frequently go extinct, and many factors such as those contributing to extinction vortices have been promoted as mechanisms leading to their decline and extinction. The genetic factors limiting the viability of small populations are believed to hinge on declining levels of variation (Frankham 2005; Allendorf et al. 2013). This reduction in variation is anticipated because the role of genetic drift should intensify as populations shrink (Allendorf 1986). Lower levels of genetic variation imply that populations will contain less of the necessary heritable variation to adapt to changes in their environment, and a larger portion of the genetic load should become fixed due to chance (Willi et al. 2006; Dussex et al. 2023). However, many island taxa persist with restricted population sizes, and their circumstances suggest they can do so for thousands of years (Frankham 1997, 1998). The populations of *Cyclura carinata* studied here have likely experienced isolation since sea levels rose at the onset of the current interglacial period and partially inundated the island banks where they reside (Welch et al. 2017). Given that these populations have been restricted by island size, they provide an opportunity to explore potential genetic factors that might facilitate or limit their viability. For example, gene flow among populations, or metapopulation dynamics, might bolster effective population sizes realized by island populations (Turnock et al. 2025). It has also been argued that taxa can evolve resistance to small population size and inbreeding depression (Robinson et al. 2018). Perhaps inbreeding avoidance mechanisms have evolved that facilitate the maintenance of genetic variation greater than that predicted by genetic drift alone (Keller and Waller 2002b). It is conceivable that *C. carinata* owes its persistence on small cays of the Turks and Caicos Islands to a combination of selection against homozygous individuals (genetic purging) and metapopulation dynamics countering the loss of genetic variability via drift.

There is limited evidence for ongoing gene flow between the *C. carinata* study populations. We found mixed results suggesting that the two regions of sampled cays are operating as indepedent metapopulations. Pairwise *F_ST_* values were between 0.11 and 0.39 within Chalk Sound National Park and within the outlying cays to the northeast, comparable to the global *F_ST_*= 0.16 found across the western lineage in Welch et al. 2017. These moderate values of *F_ST_*indicate restricted gene flow, as also evident by the region’s water barriers between cays, which does not support metapopulation dynamics.

On the other hand, our DAPC and STRUCTURE analyses show minimal cay subpopulation differentiation. Chalk Sound National Park samples are best described with genetic cluster K=1 with cay overlap in Fig. 5A. This provides evidence contrary to the pairwise *F_ST_* values that Chalk Sound National Park cays are functioning as a metapopulation. This is within the realm of possibility as Chalk Sound National Park consists of a shallow lagoon with weak tide dynamics, creating ideal conditions for iguanas to raft or swim between islands. These protected cays are also extremely small with limited carrying capacities, which may lead to greater pressure to disperse to other cays in response to competition (Case and Bolger 1991). Another explanation for the patterns seen is ancestral migration from the nearby, large island of Providenciales to Chalk Sound. This mainland-island colonization hypothesis would also result in seemingly panmictic genetic signatures. The iguana population on Providenciales has been extirpated since at least 1970 (Iverson 1979), thereby cutting off a source population for Chalk Sound. This could lead to eventual local extinction in the national park if these cay subpopulations are dependent on migrants from Providenciales. Iguanas are still present on these sampled cays as of 2015, almost twenty years after this study’s genetic samples were collected (Gerber pers. comm.).

The outlying cays northwest of Chalk Sound National Park also show minimal cay subpopulation differentiation, but more structure than the cays within the national park. Pine Cay separates out at small values of K (K=4) and does not overlap with other cays (Fig. 4), indicating uniqueness from all other cays sampled. Welch et al. 2017 collected only one sample from Pine Cay and thus were left with inconclusive population structure results. They posited that Pine Cay seems genetically intermediate between the western and eastern ESUs, which was hypothesized based on coloration and morphometrics (Iverson 1979). Based on our data, Pine Cay had the highest genetic diversity, even with low sample size. Though we only used microsatellites in this study, Pine Cay does appear genetically intermediate, sharing some but not all genetic signatures with the western ESU.

K=2 is the best fit for describing genetic clustering for the rest of the outlying cays, which consistently grouped CILC with CIBC and CIDC, CIMG, and CILW together. Fig. 5B shows separation even between CILC and CIBC, which is surprising given they are only separated by a ∼130m long channel. While short distances separate these outlying cays, the tide dynamics and currents of these channels are powerful (Chérubin 2014). Migrant iguanas may be more likely swept out to sea than successfully establish on a new cay, limiting gene flow between these subpopulations. We feel this evidence provides only weak support for metapopulation dynamics among these outlying cays.

Despite the potential for low rates of gene flow among islands, there is clear evidence of inbreeding and inbreeding depression within island populations. This is well supported by the positive correlations among genetic diversities, inbreeding coefficients, and island sizes. As the sampled island area increased, expected heterozygosity also increased. In general, more island area means more iguana habitat and thus a larger iguana population. It is typical for larger populations to have greater amounts of genetic variation (Wright 1969, Allendorf et al. 2013). Oddly, inbreeding coefficients (*F_IS_* values) also increase with island size. Paradoxically, this implies that a greater number of individuals in larger populations are inbred relative to that observed in smaller populations. One explanation for this finding is that homozygosity is better correlated with inbreeding in smaller populations. In such a case, selection associated with inbreeding depression would more readily favor heterozygotes in smaller populations. *F_IS_* estimates from larger islands in the outlying cays were larger relative to those of the smaller islands in Chalk Sound National Park. Since homozygosity is more likely to correlate with inbreeding, this may simply be a signature of soft selection acting against more homozygous individuals (Christiansen 1975; Whitlock 2002). This has been presented in other rock iguana studies (Berk 2013; Colosimo 2016; Moss et al. 2019) (but see Colosimo et al. 2026).

In addition to the evidence for elevated inbreeding depression on smaller cays, our measures of *N_E_* correlated only weakly with island size. These estimates seem largely inconsistent with our expectations given the population size estimates based on available habitat. The estimates of *N_E_* most consistent are those based simply on expected heterozygosity (Crow and Kimura 1970). However, these estimates of effective population size, while correlated with our habitat area-based estimates, are low for larger populations and high for smaller populations. This could be a function of the mutation rate used in these estimates, but using an alternate value would exacerbate differences in estimates on one end of the scale or the other, as we already use a conservative value for (Sun et al. 2012). Another factor that might bolster effective population size in the smaller populations is gene flow, and while pairwise *F_ST_* values imply low gene flow, it is conceivable that rare migration events have lasting impacts on these smaller islands (Steinbach et al. 2018). The counter argument here is that we do not see elevated estimates of effective population size for the larger populations in the study.

One assumption of these methods for estimating effective population size that is likely violated is that of no selection (Charlesworth 2009). At one level, the general theory designed to explain reduced population viability in small populations accounts for selection; purging of deleterious alleles is dependent on selection. However, strong selection at some loci should result in reduced effective population size throughout the rest of the genome, increasing the chances of fixing more mildly deleterious alleles (Schou et al. 2017). This creates linkage blocks that are selected for while not completely purging the genetic load (in other words, associative overdominance) (Gilligan et al. 2005). We hypothesize that the increased proportion of the genetic load exposed to selection in smaller populations may be facilitating the maintenance of genetic variation. To explore these dynamics further in this study system, genomic sequencing is required, though the fact that a few microsatellites scattered about the genome have revealed these strong signatures of selection is telling.

While confidence in our estimates of *N_E_* is low, it is worth noting that *N_E_* generally increased with island size (Table 3). The small values of *N_E_* match expectations with *C. carinata* habitat needs and territoriality (30 adults per hectare). Most sampled cays are very small, thereby only able to support a few iguanas at a time. We acknowledge that our analyses may be impacted by small sample size, especially with calculations of *F_IS_* and *N_E_* (Waples and Do 2008; Do et al. 2014). However, our sampling methods may have captured most iguanas on the smaller cays, thus making small sample sizes unavoidable. The assumption of a direct correlation between island size and habitat quantity may be violated in our study system without a meaningful quantification of desirable iguana habitat. For example, Mangrove Cay, as its name suggests, is composed largely of mudflats and Salinas habitat full of mangroves, while *Cyclura carinata* prefer rocky and sandy substrate amongst sparse woody vegetation (Gerber et al. 2020).

*C. carinata* may be more adept at functioning in small population sizes through adaptations for inbreeding and purging the genetic load via natural selection. Island systems can especially heighten this in vertebrates, as there is often slower (fewer instances of first order relatives mating) and consistent levels of inbreeding that decrease genetic load (Reed et al. 2003). In a *Peromyscus* study that compared species with different population histories, including mainland origin vs barrier island origin, the island species was less impacted by inbreeding in a laboratory setting than its sister taxa (Lacy and Ballou 1998). However, the population had lower vital rates before the experiment than the others. Whole genome sequencing has also confirmed effective purging of the genetic load, resulting in population recovery in endangered island species like island foxes, the Seychelles paradise flycatcher, and the red-headed wood pigeon (Robinson et al. 2018; Femerling et al. 2023; Tsujimoto et al. 2025) (but see Kennedy et al. 2014). There is a trade-off, however, between purging and decreasing genetic diversity, which can have long-term impacts on adaptation and evolutionary potential (Barthe et al. 2022). This is especially concerning with climate change and its impacts on *C. carinata*, such as rising sea levels, more major weather events, and changes in precipitation (Sealey et al. 2019).

Though not as genetically diverse as the robust CILW subpopulation, we argue that the iguana-inhabited small cays surrounding Providenciales are not currently facing extirpation. These genetic differences from CILW can be attributed to changes in allele frequencies via genetic drift, especially given the fact that these subpopulations have been established and isolated for thousands of generations (Welch et al. 2017). The differences are not severe enough to warrant conservation concern and human intervention at this time, such as assisted migration or other forms of conservation translocations.

Our findings have direct impacts on the planning of future conservation translocations of *C. carinata* and similar species. Currently, six conservation translocations have occurred in the TCI with more planned in the coming years (Gerber et al. 2007). Projects show repeatedly that successful conservation translocations are directly related to the number of individuals moved (Jamieson 2011; White et al. 2018, 2020). It is commonly recommended to have large founding populations to maximize future effective population size and curb impacts from demographic and environmental stochasticity (IUCN/SSC 2013). Supplementation after initial translocations often follows to increase the new population’s chance of long-term viability and adaptability (White et al. 2020). However, there can be many obstacles to translocating many organisms over multiple years, such as limited resources (e.g. money, personnel, equipment, permits), negative impacts on the source population, and short time frames (Bubac et al. 2019; Easton et al. 2020). The results of this study suggest empirically that populations of *C. carinata* can sustain themselves with as few as 15 individuals, at least in the short-term. This is best portrayed by Lizard and Bird Cays, two isolated cays <0.5ha with iguana subpopulations showing ample genetic diversity, limited inbreeding, and population differentiation. However, we stress that our observations of very small *C. carinata* census sizes persisting over long timescales are not the same as estimating a formal minimum viable population size or conducting a population viability analysis. As the use of conservation translocations continues to increase to prevent species extinction, harnessing successful founding population dynamics is essential for long-term endangered species management and preservation of biodiversity.

## Acknowledgements

We thank Arash Bissell, George Waters, Joe Burgess, and Gabriele Colosimo for assistance with fieldwork, and Blaklie Mitchell and Jordan Walters for laboratory assistance. We also thank the Turks and Caicos Department of Environment and Coastal Resources (DECR) for providing research permits and CITES export permits, and the IUCN/SSC Iguana Specialist Group for arranging CITES import permits. Logistical assistance in the Turks and Caicos Islands was provided by Ken Wiley, Ethlyn Gibbs-Williams, Sarah Havery, Mark Woodring, the DECR, and the Marina at South Bank. The Pine Cay Homeowners Association and the island’s general managers, Christian and Sandrine Langlade, generously provided access to their island, transportation on Pine Cay, and sustenance for our field team. Samples obtained in 2020 were collected using methods approved by the Zoological Society of San Diego Institutional Animal Care and Use Committee (IACUC), NIH assurance A3675-01.

## Data Availability

We have deposited the primary data, including sampling locations and microsatellite genotypes, underlying these analyses in Dryad. R script is also available upon request.

