## Supplemental Material for "How small is too small? Genetic signatures from naturally small populations of a large island vertebrate, the Turks and Caicos Rock Iguana"

**Supplemental Table 1:** The 28 microsatellite loci used in this study and original publication source, number of alleles found in 159 iguanas (*A*), mean observed heterozygosity (*H_O_*), mean expected heterozygosity (*H_E_*), and mean *F_IS_* (Weir and Cockerham 1984) calculated in package adegenet. Asterisks designate loci that were monomorphic and not used in downstream analyses.

| *Marker* | *Source* | *A* | *Mean H_O_* | *Mean H_E_* | *Mean F_IS_* |
| --- | --- | --- | --- | --- | --- |
| Z151 | An et al. 2004 | 5 | 0.182 | 0.584 | 0.689 |
| Z494 | An et al. 2004 | 7 | 0.080 | 0.596 | 0.865 |
| Ccste_02 | Rosas et al. 2008 | 4 | 0.139 | 0.523 | 0.734 |
| Ccste_04 | Rosas et al. 2008 | 5 | 0.559 | 0.429 | -0.301 |
| Ccste_05 | Rosas et al. 2008 | 6 | 0.425 | 0.724 | 0.413 |
| Ccste_76 | Rosas et al. 2008 | 4 | 0.336 | 0.523 | 0.358 |
| C6 | Lau et al. 2009 | 4 | 0.274 | 0.476 | 0.425 |
| C103 | Lau et al. 2009 | 6 | 0.222 | 0.360 | 0.384 |
| C113 | Lau et al. 2009 | 5 | 0.378 | 0.538 | 0.297 |
| D1 | Lau et al. 2009 | 5 | 0.281 | 0.575 | 0.510 |
| D9 | Lau et al. 2009 | 11 | 0.296 | 0.596 | 0.503 |
| D11 | Lau et al. 2009 | 7 | 0.312 | 0.584 | 0.466 |
| D107 | Lau et al. 2009 | 4 | 0.335 | 0.430 | 0.220 |
| D110 | Lau et al. 2009 | 5 | 0.301 | 0.453 | 0.336 |
| D111 | Lau et al. 2009 | 7 | 0.510 | 0.683 | 0.254 |
| D130 | Lau et al. 2009 | 3 | 0.110 | 0.195 | 0.437 |
| D137 | Lau et al. 2009 | 5 | 0.410 | 0.540 | 0.240 |
| D140 | Lau et al. 2009 | 6 | 0.446 | 0.660 | 0.324 |
| CIDK101 | Welch et al. 2011 | 3 | 0.110 | 0.167 | 0.338 |
| *CIDK109 | Welch et al. 2011 | 1 | 0.000 | 0.000 | 0.000 |
| CIDK113 | Welch et al. 2011 | 3 | 0.123 | 0.536 | 0.770 |
| *CIDK135 | Welch et al. 2011 | 1 | 0.000 | 0.000 | 0.000 |
| CIDK144 | Welch et al. 2011 | 4 | 0.229 | 0.257 | 0.109 |
| CIDK163 | Welch et al. 2011 | 4 | 0.342 | 0.555 | 0.384 |
| CIDK177 | Welch et al. 2011 | 4 | 0.284 | 0.426 | 0.334 |
| CIDK184 | Welch et al. 2011 | 2 | 0.139 | 0.249 | 0.440 |
| CIDK199 | Welch et al. 2011 | 2 | 0.131 | 0.353 | 0.630 |
| CIDK228 | Welch et al. 2011 | 2 | 0.085 | 0.081 | -0.044 |

**Supplemental Table 2:** Table of the Evanno method output for all sampled cays, K=1-13. Values in bold are for the subpopulation with the largest Delta K, representing primary genetic clustering.

| *K* | *nRep* | *Mean LnP (K)* | *Stdev LnP (K)* | *Ln’ (K)* | *\|Ln’’ (K)\|* | *Delta K* |
| --- | --- | --- | --- | --- | --- | --- |
| 1 | 20 | -6616.58 | 10.92 | NA | NA | NA |
| **2** | **20** | **-5326.21** | **8.54** | **1290.37** | **870.84** | **101.972** |
| 3 | 20 | -4906.68 | 25.21 | 419.53 | 246.54 | 9.779 |
| 4 | 20 | -4733.69 | 209.04 | 172.99 | 36.52 | 0.175 |
| 5 | 20 | -4524.18 | 85.99 | 209.51 | 29.94 | 0.348 |
| 6 | 20 | -4344.61 | 72.41 | 179.57 | 32.46 | 0.448 |
| 7 | 20 | -4197.5 | 49.34 | 147.11 | 46.31 | 0.939 |
| 8 | 20 | -4096.7 | 61 | 100.8 | 97.13 | 1.592 |
| 9 | 20 | -4093.03 | 68.14 | 3.67 | 74.88 | 1.099 |
| 10 | 20 | -4014.48 | 97.72 | 78.55 | 16.31 | 0.167 |
| 11 | 20 | -3919.62 | 95.15 | 94.86 | 47.61 | 0.5 |
| 12 | 20 | -3872.37 | 81.47 | 47.25 | 14.12 | 0.173 |
| 13 | 20 | -3811 | 46.25 | 61.37 | NA | NA |

**Supplemental Table 3:** Table of the Evanno method output for sampled cays in Chalk Sound National Park, K=1-7. Values in bold are for the subpopulation with the largest Delta K, representing primary genetic clustering.

| *K* | *nRep* | *Mean LnP (K)* | *Stdev LnP (K)* | *Ln’ (K)* | *\|Ln’’ (K)\|* | *Delta K* |
| --- | --- | --- | --- | --- | --- | --- |
| 1 | 20 | -1695.42 | 6.15 | NA | NA | NA |
| 2 | 20 | -1558.2 | 14.53 | 137.22 | 14.4 | 0.991 |
| **3** | **20** | **-1435.38** | **10.34** | **122.82** | **34.68** | **3.354** |
| 4 | 20 | -1347.24 | 18.44 | 88.14 | 31.95 | 1.733 |
| 5 | 20 | -1291.05 | 19.73 | 56.19 | 16.92 | 0.858 |
| 6 | 20 | -1251.78 | 16.98 | 39.27 | 35 | 2.061 |
| 7 | 20 | -1247.51 | 29.33 | 4.27 | NA | NA |

**Supplemental Table 4:** Table of the Evanno method output for sampled cays in the outlying cays northeast of Provo, K=1-5. Values in bold are for the subpopulation with the largest Delta K, representing primary genetic clustering.

| *K* | *nRep* | *Mean LnP (K)* | *Stdev LnP (K)* | *Ln’ (K)* | *\|Ln’’ (K)\|* | *Delta K* |
| --- | --- | --- | --- | --- | --- | --- |
| 1 | 20 | -3033.88 | 10.97 | NA | NA | NA |
| **2** | **20** | **-2634.03** | **10.71** | **399.85** | **240.46** | **22.452** |
| 3 | 20 | -2474.64 | 14.28 | 159.39 | 86.25 | 6.04 |
| 4 | 20 | -2401.5 | 59.9 | 73.14 | 47.23 | 0.788 |
| 5 | 20 | -2281.13 | 20.25 | 120.37 | NA | NA |

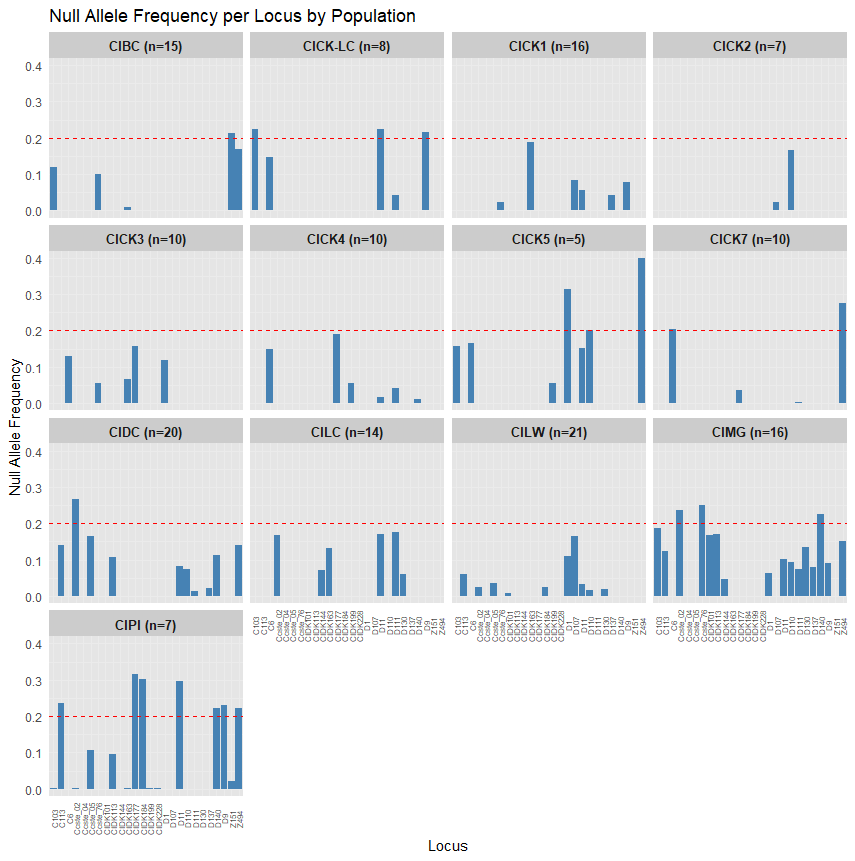

**Supplemental Figure 1:** Null allele frequency per locus by sampling location as calculated using the Expectation Maximization algorithm method in the program FreeNA. The red dashed line at 0.20 indicates the lower threshold for presence of null alleles.

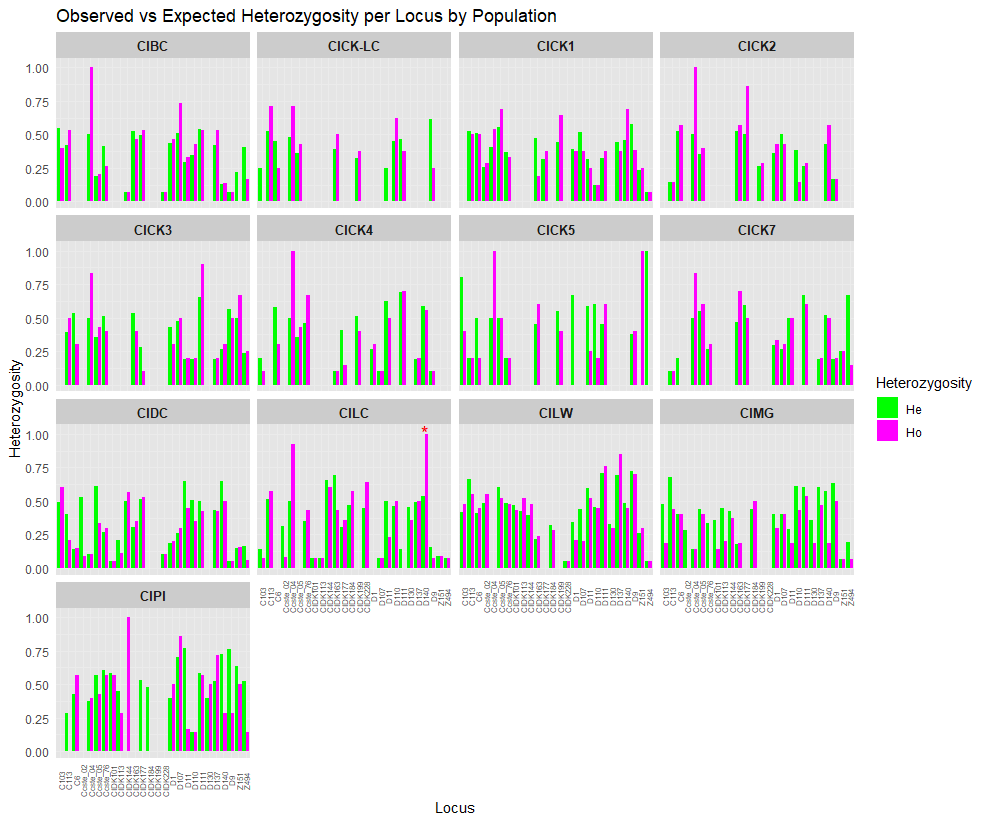

**Supplemental Figure 2:** Observed (Ho) vs expected heterozygosity (He) per locus by sampling location used to check for Hardy-Wienberg equilibrium. Red asterisks depict significant differences based on an exact test and corrected for false discovery rate.

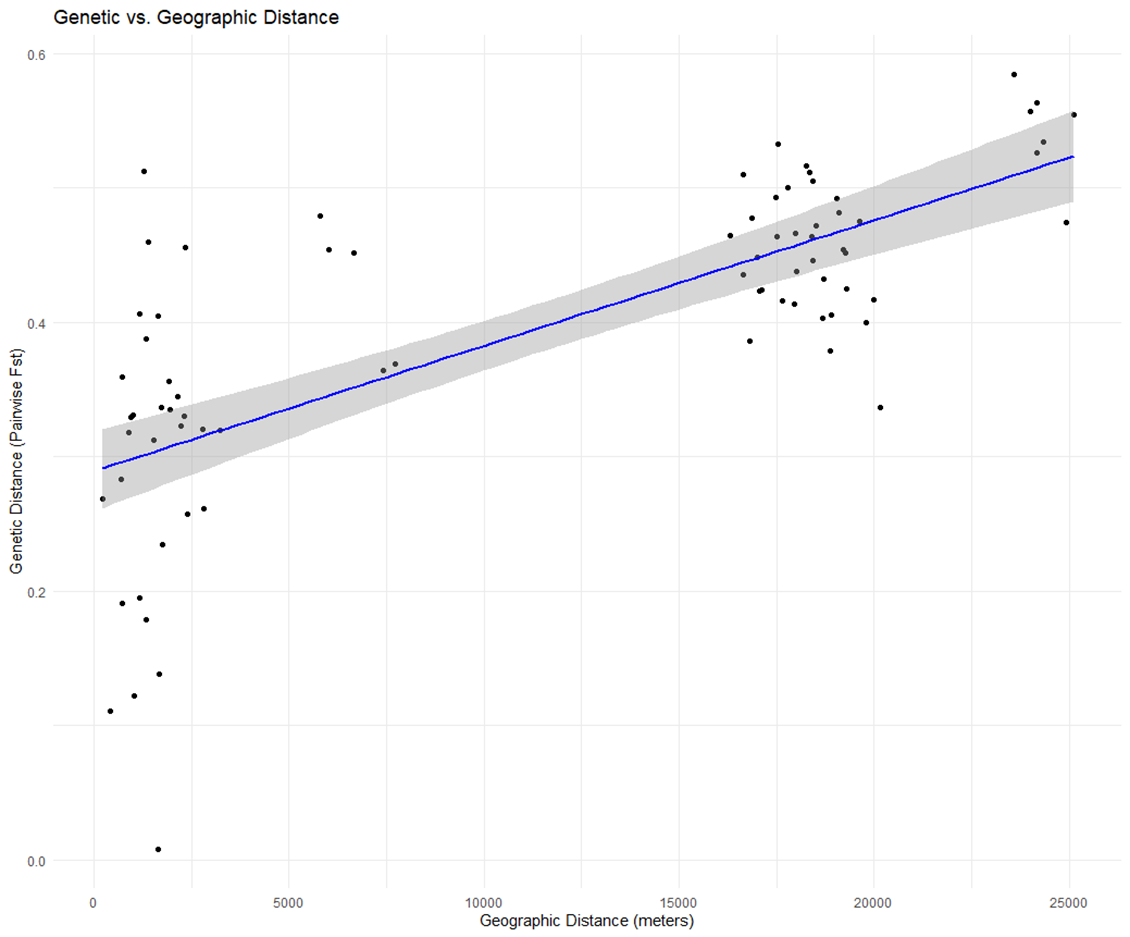

**A**

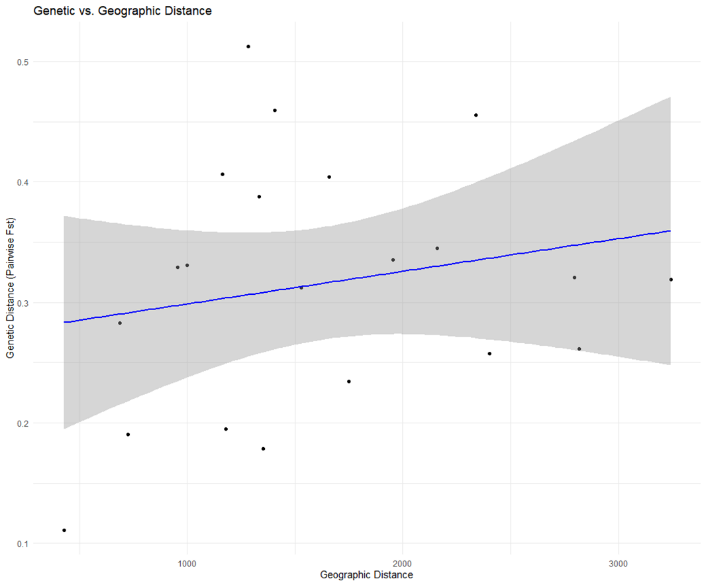

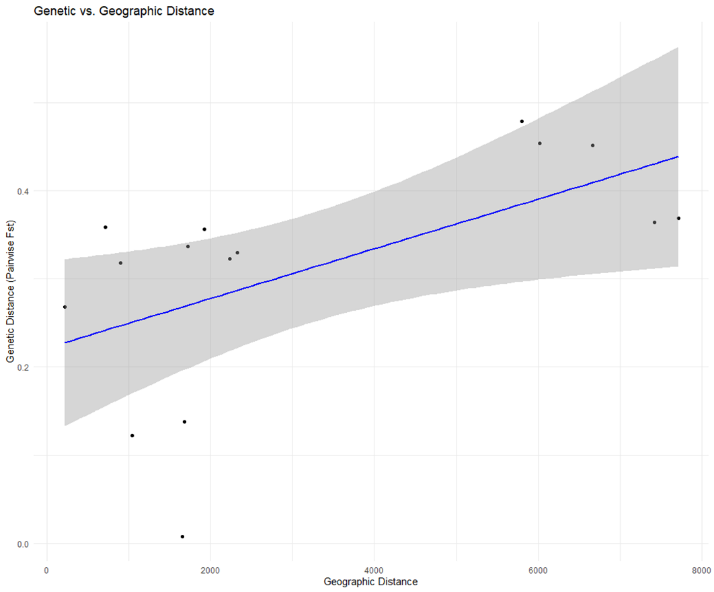

**C**

**B**

**Supplemental Figure 3:** Isolation by distance model for A) all cays, B) Chalk Sound National Park and C) outlying cays. Geographic distance is in meters and genetic distance is represented using Weir and Cockerham’s pairwise *F_ST_*. Shaded area is standard error.

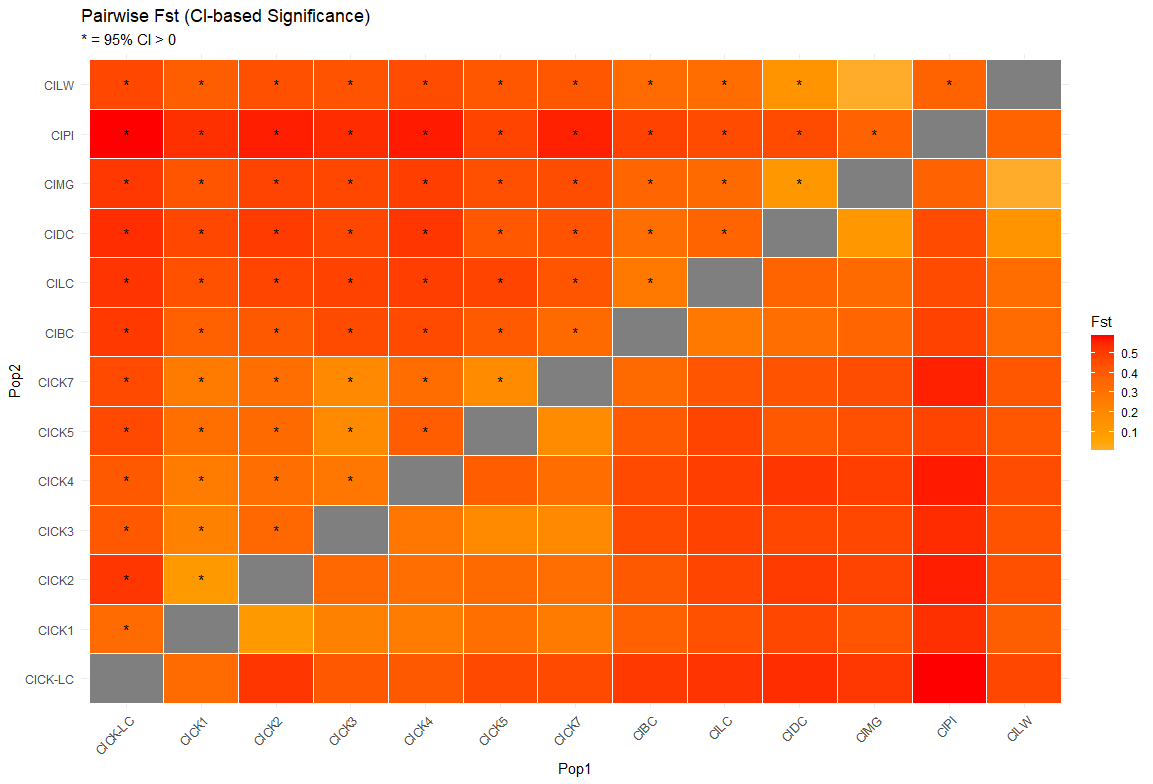

**Supplemental Figure 4:** Heat map showing pairwise *F_ST_* comparisons across all sampled cays. Asterisks indicate comparisons where the lower confidence internals show a significant difference from 0. *F_ST_* values range from 0.0076 to 0.5845.

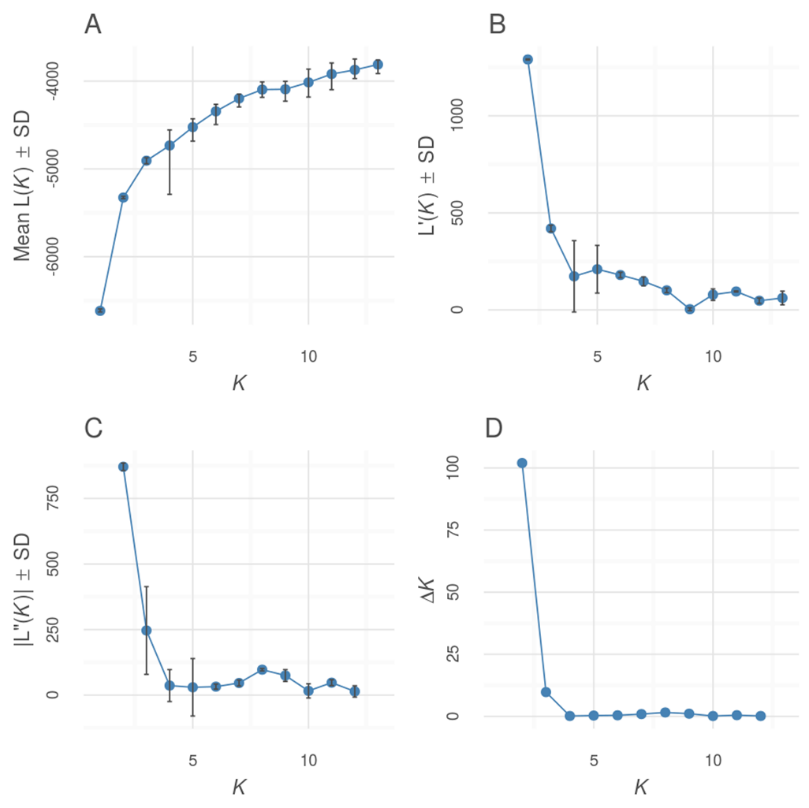
E

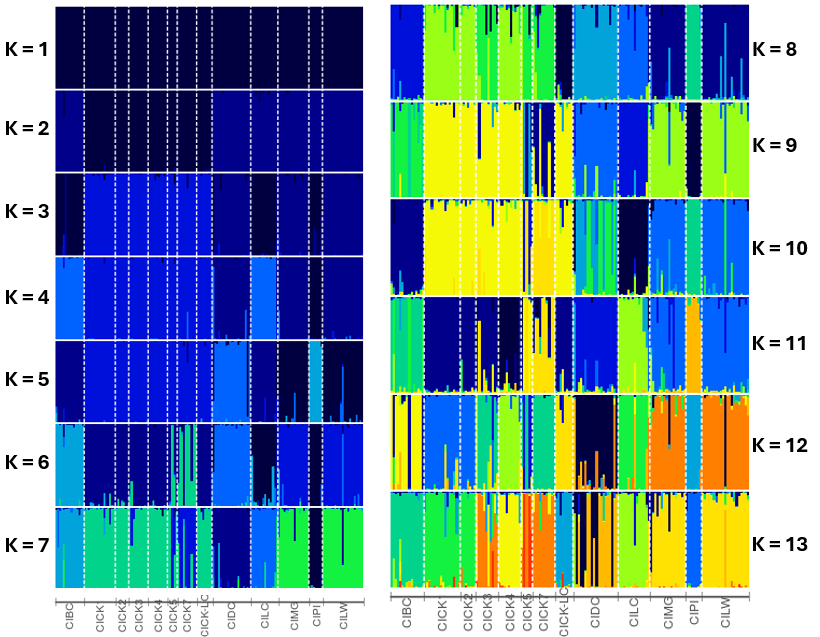

**Supplemental Figure 5:** A-D) Graphing results of the Evanno method output for all cays, K=1-13. E) Distruct plots from STRUCTURE output for all sampled cays produced using web app PopHelper. Each bar represents one iguana, each color represents best fit genetic cluster, and dotted lines separate samples by their cay.

E
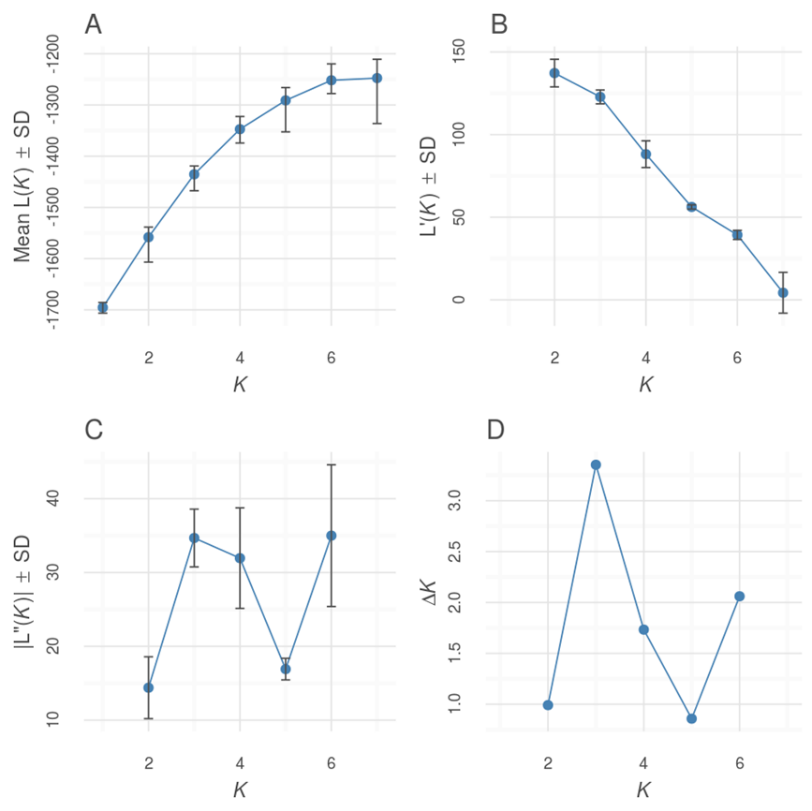

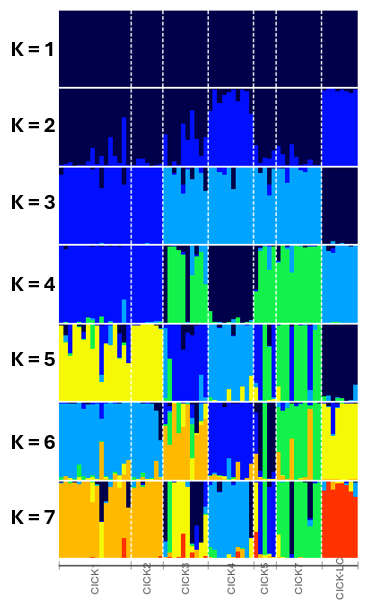

**Supplemental Figure 6:** A-D) Graphing results of the Evanno method output for sampled cays in Chalk Sound National Park, K=1-7. See Table S4 for more details. E) Distruct plots from STRUCTURE output for Chalk Sound National Park cays produced using web app PopHelper. Each bar represents one iguana, each color represents best fit genetic cluster, and dotted lines separate samples by their cay.

E
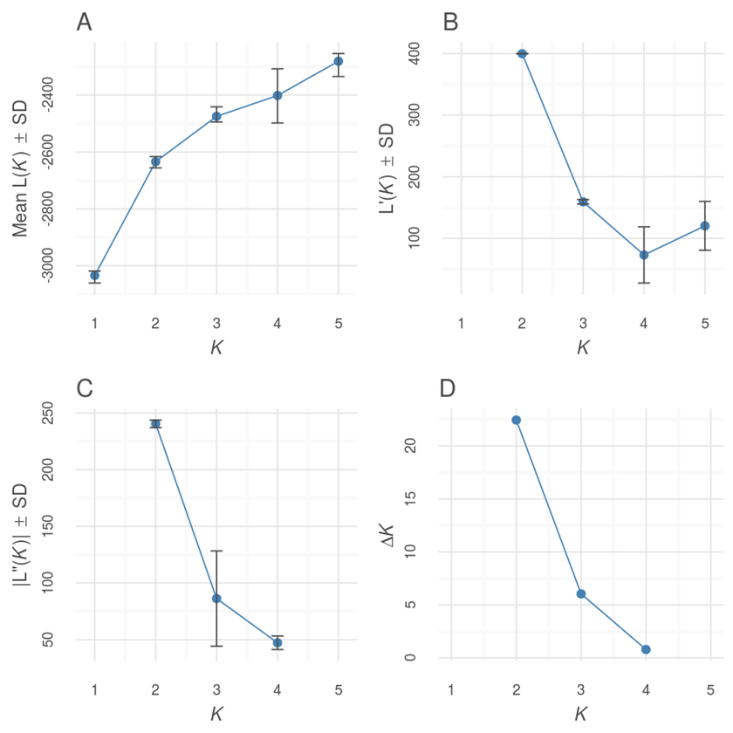

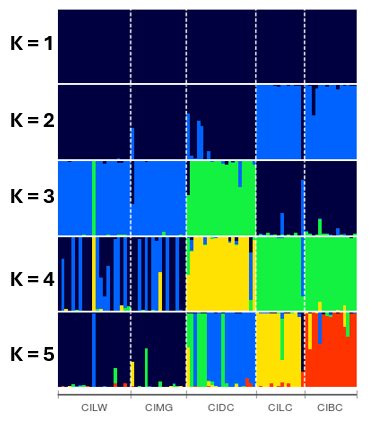

**Supplemental Figure 7:** A-D) Graphing results of the Evanno method output for sampled cays northeast of Provo, K=1-5. See Table S5 for more details. E) Distruct plots from STRUCTURE output for the outlying cays northeast of Provo produced using web app PopHelper. Each bar represents one iguana, each color represents best fit genetic cluster, and dotted lines separate samples by their cay.
